# Self-Supervised Contrastive Learning of Protein Representations By Mutual Information Maximization

**DOI:** 10.1101/2020.09.04.283929

**Authors:** Amy X. Lu, Haoran Zhang, Marzyeh Ghassemi, Alan Moses

**Affiliations:** insitro; Department of Computer Science, University of Toronto; Vector Institute for Artificial Intelligence; Department of Cell and Systems Biology, University of Toronto

## Abstract

Pretrained embedding representations of biological sequences which capture meaningful properties can alleviate many problems associated with supervised learning in biology. We apply the principle of mutual information maximization between local and global information as a self-supervised pretraining signal for protein embeddings. To do so, we divide protein sequences into fixed size fragments, and train an autoregressive model to distinguish between subsequent fragments from the same protein and fragments from random proteins. Our model, CPCProt, achieves comparable performance to state-of-the-art self-supervised models for protein sequence embeddings on various downstream tasks, but reduces the number of parameters down to 2% to 10% of benchmarked models. Further, we explore how downstream assessment protocols affect embedding evaluation, and the effect of contrastive learning hyperparameters on empirical performance. We hope that these results will inform the development of contrastive learning methods in protein biology and other modalities.

## 1. Introduction

Due to improved sequencing technologies, the size of protein databases have seen exponential growth over the past decades (Consortium, 2019). However, the cost and time associated with obtaining labels for *supervised* problems on these proteins presents an important challenge. In response, recent works look towards self-supervised pretraining for obtaining informative fixed-length embeddings from amino acid sequences. Given input sequences, a loss is minimized that does not rely on labels related to the quantity of interest (e.g. function or structure), but derived from the data itself. This circumvents the need to obtain experimental or expert-annotated labels.

Self-supervised learning based on contrastive tasks aim simply to tell apart aspects of the data, as opposed to generative tasks, such as inpainting (Pathak et al., 2016) or autoencoding (Kingma & Welling, 2013), which train models to generate part or all of the data. A major advantage of contrastive learning, in principle, is that no complicated decoding of the latent space is required (Oord et al., 2018). Moving model capacity to the encoder to obtain embeddings with high predictive power is appealing for several reasons. First, they can be portable for computational biologists without the computational resources to train large models. Second, collapsing sequence data into an informative latent space help with clustering and leveraging similarity information between proteins to learn more about protein mechanisms and function.

Recent works demonstrate self-supervised pretraining of protein sequences can yield embeddings which implicitly capture properties such as phylogenetic, fluorescence, pair-wise contact, structural, and subcellular localization. However, many of these works directly take embedding techniques from natural language processing (NLP) tasks (Yang et al., 2018a; Riesselman et al., 2019; Rives et al., 2019; Alley et al., 2019; Heinzinger et al., 2019; Elnaggar et al., 2019; Armenteros et al., 2019; Madani et al., 2020; Elnaggar et al., 2020). Presumably, using a more biologically-motivated proxy task will yield better insights and performance on biological data. Some methods incorporate biological information such as protein-protein interactions (Nourani et al., 2020), or structured labels from SCOP (Bepler & Berger, 2019) and PDB (Gligorijevic et al., 2019); however, high-quality curation of these labels circle back to the need for expensive experiments.

Recently, there has been growing interest in using a mutual information (MI) maximization objective for obtaining self-supervised representations (Tschannen et al., 2019). At its core, biological sequences prescribe a set of information for function, structure, etc.; the ideas of information content and error-correcting codes seem therefore to be natural fits for modelling the evolutionary and functional constraints which drove protein sequences towards its currently observed complexity. Indeed, information theoretic principles have been explored in computational biology since the the 1970s (Gatlin et al., 1972), and subsequently applied to many modelling problems in biological sequence analysis (Roman-Roldan et al., 1996; Vinga, 2014) and molecular biology (Adami, 2004), such as sequence logo visualization (Schneider & Stephens, 1990), transcription factor binding site discovery (Stormo, 2000), structure prediction of protein loops (Korber et al., 1993), and evolutionary conservation of sequence features (Rao et al., 1979; Pritišanac et al., 2019). Using the information content of protein sequence patches as a pretraining objective better aligns with underlying biological principles as compared to directly lifting methods from NLP to biology. If patches capture motifs, unusual structural elements, regions of unusual amino acid composition, parts of catalytic sites, etc., the model must capture this implicit information in its latent representation of the protein during self-supervised pretraining.

In this work, we present CPCProt, which maximize mutual information between context and local embeddings by minimizing a contrastive loss. On protein-related downstream benchmarks (Rao et al., 2019), CPCProt achieves comparable results, despite using 2% the number of pretraining parameters of the largest model (Rao et al., 2019) and 10% of the number of parameters of the smallest neural network model (Alley et al., 2019), as illustrated in Figure 1. We further explore difficulties in using downstream performance as a means to assess protein embeddings, and explore the effect of contrastive learning parameters, to motivate the development of similar methods in protein biology and other modalities.

**Figure 1.**
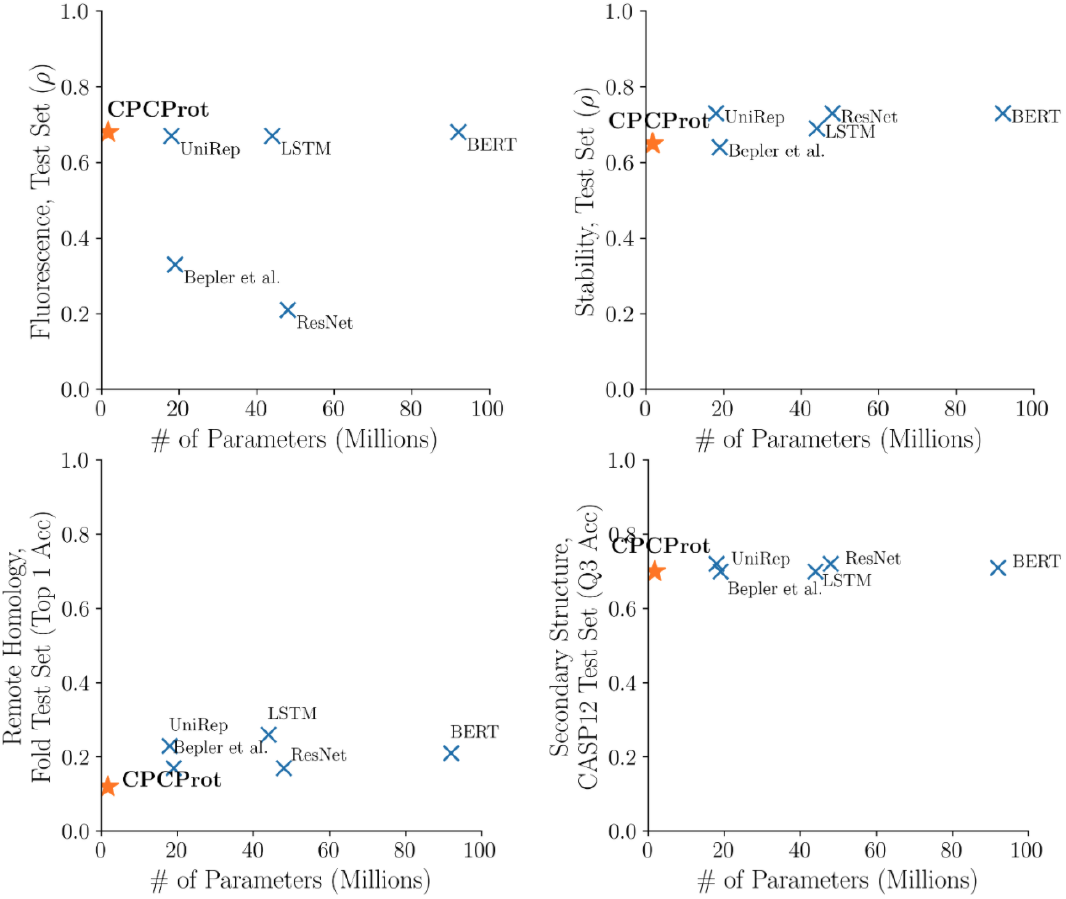
Downstream performance on protein-related tasks obtained by finetuning embeddings from pretrained models, plotted against the number of parameters in the pretrained model. Orange stars denote our model, and blue crosses denote methods bench-marked in Rao et al. (2019). *ρ* denotes Spearman’s correlation for regression tasks, and “Acc” indicates accuracy.

## 2. Background and Related Work

The InfoMax optimization principle aims to find a mapping *g* (constrained by a function class 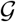), which maps from input *X* to output *g*(*X*), such that the Shannon mutual information between the pair is maximized (Linsker, 1988):

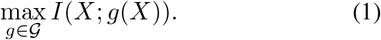

Conceptually, therefore, the training task of finding a desirable encoder which maps input sequences to output embeddings is performing InfoMax optimization.

It is possible to use a variational approach to estimate a bound on the mutual information between continuous, high-dimensional quantities (Donsker & Varadhan, 1983; Nguyen et al., 2010; Alemi et al., 2016; Belghazi et al., 2018; Oord et al., 2018; Poole et al., 2019). Recent works capture this intuition to yield self-supervised embeddings in the modalities of imaging (Oord et al., 2018; Hjelm et al., 2018; Bachman et al., 2019; Tian et al., 2019; Hénaff et al., 2019; Löwe et al., 2019; He et al., 2019; Chen et al., 2020; Tian et al., 2020; Wang & Isola, 2020), text (Rivière et al., 2020; Oord et al., 2018; Kong et al., 2019), and audio (Löwe et al., 2019; Oord et al., 2018), with high empirical downstream performance.

The general formulation is: given input *X*, define {*X*^(1)^*, X*^(2)^} as two different “views” of *X* (e.g. patches of an image, or representations of different sequence timesteps), and encoders {*g*_1_*, g*_2_} which encode {*X*^(1)^*, X*^(2)^} respectively. The goal is to find encoder mappings which maximize the mutual information between the outputs, and can be shown to lower bound the InfoMax objective in Equation 1 (Tschannen et al., 2019):

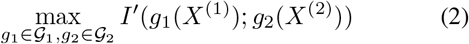

## 3. Methods

We describe CPCProt, which applies the InfoNCE loss introduced by the Contrastive Predictive Coding (CPC) method (Oord et al., 2018) to protein sequences. Pretrained model weights are available at hershey.csb.utoronto.ca/CPCprot/weights/. Code for embedding single sequences and FASTA files, as well as evaluation code for reproducibility is at github.com/amyxlu/CPCProt.

### 3.1 Contrastive Predictive Coding and InfoNCE

Here, we formalize the InfoNCE loss for mutual information maximization in the language of protein sequences. The CPC method introduces a lower-bound estimator for the unnormalized mutual information between two continuous quantities. It is named as such as it stems from the noise-contrastive estimation (NCE) method (Gutmann & Hyvärinen, 2010). NCE fits logistic regression parameters to distinguish data from noise (i.e. contrastively), in order to parameterize models in high dimensions, and InfoNCE directly adapts this contrastive task to estimate mutual information instead.

Define *g*_1_ and *g*_2_ from Equation 2 to be an encoder and autoregressor respectively, which we denote as *g*_*enc*_ and *g*_*ar*_. Further, define *x* as an input protein sequence, *z* as the latent embedding produced by *g*_*enc*_(*x*), and *c* as the long-range protein sequence “context”, as summarized by the autoregressor *g*_*ar*_(*z*). At a given position *t* (indexed in the *latent* space, i.e. for *z*), we compute the InfoNCE mutual information estimation 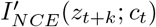 for *k* ∈ {1, 2,…, *K*}, for a batch of *N* samples. The estimation is performed by minimizing the loss:

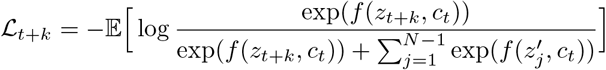

In other words, in each batch of *N* samples, we have a single sample *z*_*t*+*k*_ drawn jointly with *c*_*t*_ from *p*(*z*_*t+k*_, *c*_*t*_). Then, following the NCE method, we draw *N* − 1 “fake” samples from the noise distribution *p*(*z*′) to create a set of 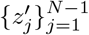. In practice, the expectation is taken over multiple batches.

This objective is a contrastive task, using a cross-entropy loss which encourages a critic, *f*, to correctly identify the single “real” sample of *z*_*t*+*k*_. Minimizing this loss provides an unnormalized lower-bound estimate on the true MI, 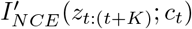 (Oord et al., 2018).

From the representation learning perspective, we obtain embeddings which maximize the mutual information between the protein “context” (e.g. long-range contacts) and the local embedding space (e.g. local secondary structure). Next-token prediction as a self-supervised proxy task has previously been successfully used to learn representations of proteins (Alley et al., 2019), and both methods share the underlying intuition that capturing latent information between current and future positions in the protein sequence may richly capture a down-sampled or “patched” version of the input amino acid sequence.

### 3.2. CPCProt

We adapt a strategy from image experiments (Oord et al., 2018; Hénaff et al., 2019; Hjelm et al., 2018; Bachman et al., 2019) to protein sequences, in which the input *x* (zero-padded up to the longest sequence in a batch) is divided into fixed-length patches. Each patch is passed into an encoder to output a single pooled embedding for the patch. These embeddings are then concatenated into the latent embedding *z*; that is, the length of *z* becomes 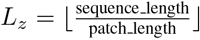. Here, a patch length of 11 is selected, such that it is long enough to capture some local structural information and also results in a reasonable *L*_*z*_ on our pretraining dataset. We start with some *t*_min_ to allow *c*_*t*_ to gain some degree of sequence context in calculations of the loss. We then calculate 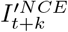 for every *t* ∈ {*t*_min_, *t*_min_ + 1,…, *L*_*z*_ − *K*} and *k* ∈ {1, 2,…, *K*}. Patching reduces the number of InfoNCE calculations necessary), and ensures that the inputs used to create *z*_1_, *z*_2_,…, *z*_*k*_ do not overlap, to reduce triviality of the task. A schematic detailing the method is illustrated in Figure 2.

**Figure 2.**
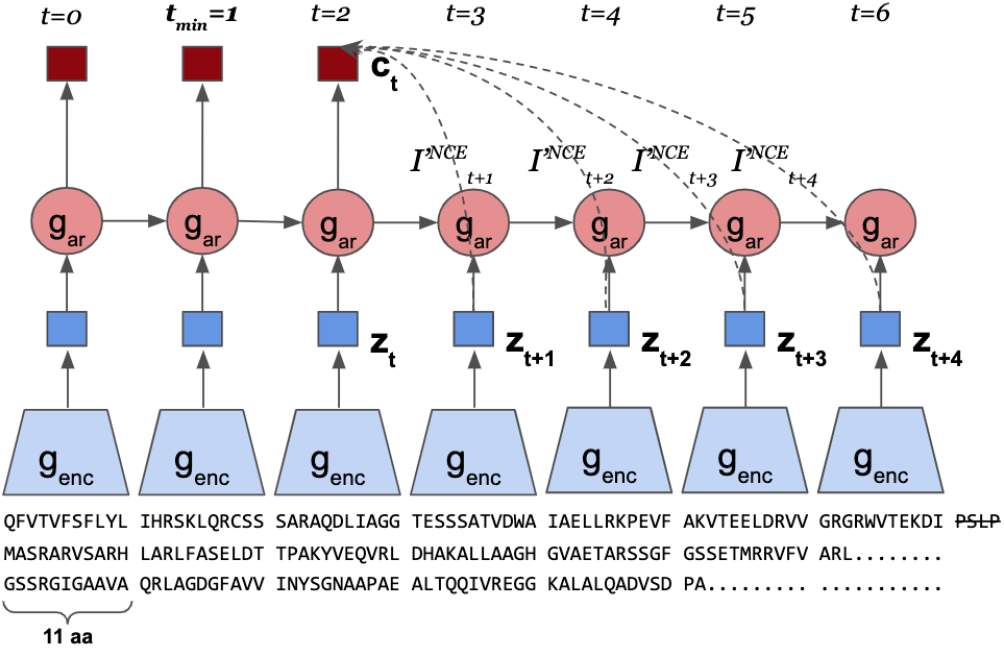
Input protein sequences are divided into “patches” of 11 amino acids. Each patch is encoded into a single vector though *g*_*enc*_, and all encodings are concatenated to form *z*. *g*_*ar*_ is an autoregressor that aggregates local information, and produce *c*, a context vector which summarizes the global context. Amino acid sequences are zero-padded to the longest sequence in the batch, with remaining amino acids not long enough for a patch discarded. For a given batch, the loss is the average of the InfoNCE estimate 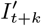 for all *t* ∈ {*t*_min_, *t*_min_ + 1,…, *L*_*z*_ − *K*} and *k* ∈ {1, 2,…, *K*}. In this example batch, *t*_min_ = 1, *L*_*z*_ = 6, and *K* = 4.

The final loss minimized in each batch is the average of calculated 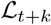 for all values of *t* and *k*, i.e.:

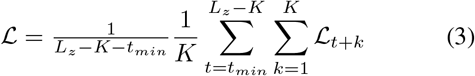

#### Architecture

In Oord et al. (2018), it is shown a simple architecture can achieve good performance with NLP tasks; we design a simple convolutional encoder with increasing number of filters in the top layers, to keep a lightweight number of connections.

The CPCProt encoder consists of embedding layer to 32 hidden dimensions, followed by 64 filters of length 4, 64 filters of length 6, and 512 filters of length 3. Filter lengths are designed such that the output of an 11 amino acid patch has an output length of one. ReLU activation is added to the output of each layer, and normalization is added channel-wise, to avoid using batch normalization, since negative samples are drawn in-batch (see Section 3.2).

In addition, we also report results using a larger encoder. Both CPCProt_GRU_large_ and CPCProt_LSTM_ uses an embedding layer to 64 hidden dimensions, followed by 128 filters of length 4, 256 filters of length 4, and 512 filters of length 3; in the final layer, CPCProt_GRU_large_ uses 1024 filters of length 3, whereas CPCProt_LSTM_ uses 2048 filters.

For both CPCProt and CPCProt_GRU_large_, a single-layer GRU is used as the autoregressor, while CPCProt_LSTM_ uses a two-layer LSTM. To avoid information leakage about later sequence locations in the context vector, we only use uni-directional autoregressors. All autoregressors use the same number of hidden dimensions as encoder output.

#### Distinguishing Noise Samples

To encourage the encoder to learn a richer embedding, and to mitigate the need to train separate critics for each position of *k*, we modify the bilinear critic used in Oord et al. (2018), and instead use a parameterless dot product critic for *f* (Chen et al., 2020). Rather than using a memory bank (Wu et al., 2018; Tian et al., 2019; He et al., 2019), we draw “fake” samples from *p*(*z*) and *p*(*c*) using other *z*_*t*+*k*_ and *c*_*t*_ from other samples in the same batch (Chen et al., 2020). That is, the diagonal of the output of the dot-product critic is the “correct pairing” of 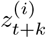 and 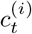 at a given *t* and *k* and the softmax is computed using all off-diagonal entries to draw the *N* −1 “fake” samples from the noise distribution *p*(*z*′).

#### Additional Pretraining Details

We use *t*_min_ = 1 and choose *K* = 4 (that is, 44 amino acids away). Sequences are zero-padded up to the longest sequence in that batch, and protein domains are truncated to a maximum of 550 amino acids due to memory constraints. Sequences shorter than the minimum length needed to perform the NCE calculation is discarded. Since *k* is 1-indexed, the minimum training sequence length is (*t*_min_ + 1 + *K*) patch length. For our default models, this minimum length is (1+1+4) × 11 = 66 amino acids. A large *K* incentives the model to pick up more information during training, but decreases computational efficiency. Furthermore, because we throw away sequences which are too short to complete the NCE loss at *t*_min_ + *K*, a large choice of *t*_min_ and *K* results in discarding more data during training.

For CPCProt, we use a batch size of 64, trained for 19 epochs with a constant learning rate of 1e-4. For CPCProt_LSTM_, a batch size of 1024 was used, trained for 18 epochs with an initial learning rate of 1e-3 and decayed by *γ* = 0.85 at each epoch. For CPCProt_GRU_large_, a batch size of 1024 is used with a constant learning rate of 1e-4, trained for 24 epochs. All models use the Adam optimizer with *β*_1_ = 0.9, *β*_2_ = 0.999, and *ϵ* = 1e-8 (Kingma & Ba, 2014). Convergence was determined as no improvement in the validation contrastive loss for 5 epochs.

## 4. Data and Downstream Evaluation

### 4.1. Pretraining

#### Pretraining Data

We pretrain our models using protein domain sequences from the Pfam database (El-Gebali et al., 2019). To compare only the effectiveness of the InfoNCE objective, we pretrain using the same data splits as used for downstream benchmarks (Rao et al., 2019), which holds out six Pfam clans as the test set. 5% of the remaining sequences are used for validation, and the remaining 32,207,059 Pfam amino acid sequences are used for pretraining.

#### Evaluating Embeddings for Hyperparameter Selection

We examine three evaluations for pretraining hyperparameter selection: (1) Performance on the validation dataset on downstream tasks (see Section 4.2); (2) contrastive accuracy on the pretraining validation data; and (3) Pfam family prediction using a 1-nearest-neighbor (1NN) classifier on the pretraining validation data. The latter two evaluation metrics are explored to avoid overfitting to benchmark datasets (Recht et al., 2019). Though in principle, the contrastive accuracy on heldout pretraining data is sufficient for hyperparameter selection, we were concerned that the contrastive task is relatively local, and may fail to assess how well embeddings have captured the global context.

The 1NN classification task is a direct measure of the ability for embeddings to cleanly separate Pfam domains in the latent space, and requires no parameter tuning or additional labels for evaluation. For this task, the dataset consists of sequences from the 50 Pfam families with the most sequences in the pretraining validation dataset, subsampled to 120 sequences per family for class balance. 70% of this embeddings is used to populate the 1NN classifier, and 30% of the sequences are used at the classification phase. A t-SNE of CPCProt embeddings colored by the 50 families is shown in Figure 3.

**Figure 3.**
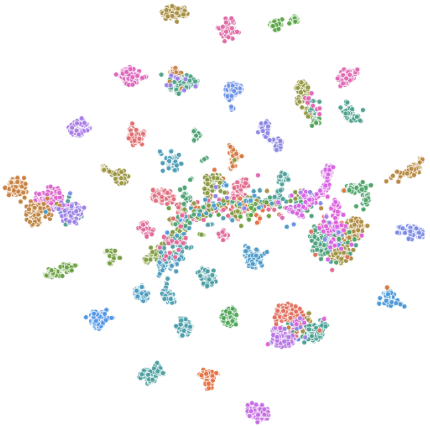
CPCProt embedding t-SNE of the 50 largest Pfam families in the validation dataset, using the final position of the context vector. Note that while colours denote different families, proximity in the continuous color space do not correspond to any intrinsic similarities between families.

For the contrastive task (i.e. the self-supervised pretraining task), we keep the ratio of negative-to-positive samples consistent and use a batch size of 512 for all models for this validation.

### 4.2. Downstream Evaluation Protocols

To understand properties of the embeddings, we examine performance on various supervised downstream tasks, derived from Tasks Assessing Protein Embeddings (TAPE) (Rao et al., 2019), using three different downstream evaluation models: a small neural network head, linear models, and k-nearest-neighbours (kNN).

#### Neural Network^1^ Finetuning Head Evaluation

For better consistency with benchmark protein-related tasks, we use the same default downstream architectures as provided by the authors in the `tape-proteins` package, version 0.4 (Rao et al., 2019). In line with benchmark methods, we allow backpropagation through the embeddings during finetuning. For consistency, we do not tune the architecture of the neural network heads. A grid search is conducted on the learning rate, batch size, and number of finetuning epochs, and test set results are reported for the model with the best performance on the validation set for a given task.

Using a non-linear neural network headwith end-to-end fine-tuning presumably yields better performance on downstream tasks, and offers a better evaluation of capabilities which can be derived from the embeddings, for those working directly with the biological tasks assessed in this work.

#### Linear and Nearest-Neighbours Downstream Evaluation

Following previous work in self-supervised learning, we also assess embeddings using a linear model (Oord et al., 2018; Hénaff et al., 2019; He et al., 2019; Bachman et al., 2019; Tian et al., 2019; 2020). Note that as compared to the neural network finetuning head evaluation, we use static embeddings extracted from the model without end-to-end optimization. For classification tasks, we use a logistic regression model, while for regression tasks, we use a linear regression model. For logistic regression models, we report the best result from conducting a simple grid search over the inverse regularization strength *C* ∈ {1, 0.01, 0.0001}.

In addition, we evaluate separation in the latent space using a kNN model with a grid search over the number of neighbours *k* ∈ {1, 5, 10}. Due to computational considerations, we do not use the kNN for secondary structure prediction.

Evaluating using a simple classifier assesses several desirable embedding properties: (1) limiting the modelling capacity given to the downstream models is a more direct assessment of the powers of the embedding methods themselves; (2) linear/logistic regression and kNN is more suitable than a neural network for downstream tasks where data availability is a constraint, as is the case for many biological use cases; (3) logistic regression has a convex loss whose optimization is relative more straight-forward than the non-convex training of a neural network. Similarly, a kNN consists simply of a look-up at test time. This arguably removes some ambiguity regarding if differences in performance should be attributed to lack of convergence or improper hyperparameter tuning when finetuning using a neural network head; and (4) as we note in Section 4.3, there exist minor architectural differences between the neural network architectures released in `tape-proteins` and the original benchmarks in Rao et al. (2019). In conjunction with the neural network head finetuning results, we hope that a more holistic picture of the embeddings can be provided.

### 4.3. Downstream Evaluation Tasks

#### Remote Homology

Remote sequence homologs share conserved structural folds but have low sequence similarity. The task is a multi-class classification problem, consisting of 1195 classes, each corresponding to a structural fold. Since global context from across the Pfam domain is important, we use the final position of the autoregressor output, *c*.

Data from the SCOP 1.75 database (Fox et al., 2013) is used. Each fold can be sub-categorized into superfamilies, and each superfamily sub-categorized into families. The training, validation, and test set splits are curated in Hou et al. (2018); test sets examines three levels of distribution shift from the training dataset. In the “Family” test set, proteins in the same fold, superfamily, and family exists in both the training and testing datasets (i.e. no distribution shift). The “Superfamily” test set holds out certain families within superfamilies, but sequences with overlap with training dataset at the superfamily level. Finally, the “Fold” test set also holds out certain superfamilies within folds. Note that severe class imbalance exists for this task, as 433 folds in the training dataset only contains one sample.

For evaluation using a neural network head, the classification architecture is a multi-layer perceptron (MLP) with one hidden layer of 512 units. Note that results in benchmarked models also train a simple dense layer to obtain an attention vector before calculating an attention-weighted mean.

#### Secondary Structure

Secondary structure is a sequence-to-sequence task evaluating the embeddings’ ability to capture local information (Rao et al., 2019). We report three-class accuracy (Q3), following the DSSP labeling system (Kabsch & Sander, 1983). Each input amino acid is mapped to one of three labels (“helix”, “strand”, or “other”), and accuracy is the percentage of correctly-labelled positions. To obtain the embedding, we use a sliding input window to obtain *z* with the same length as the input sequence, and then use *c* as the embedding to incorporate global context.

Classification results are presented on three datasets: (1) TS115, consisting of 115 protein sequences (Yang et al., 2018b); (2) CB513, consisting of 513 protein regions from 434 proteins (Cuff & Barton, 1999); and (3) free-modelling targets from the 2016 CASP12 competition, consisting of 21 protein sequences (Abriata et al., 2018; Moult et al., 2018). For training these supervised classifiers, the same validation and filtered training datasets as NetSurf-2.0 is used, where sequences with greater than 25% sequence similarity as the three test set sequences were removed from the training set (Klausen et al., 2019).

For evaluation using a neural network head, the classification architecture in `tape-proteins` is a convolutional architecture with 512 filters of size 5 and 3 in layers one and two, respectively. The original benchmarks use a higher capacity NetSurfP model (Klausen et al., 2019), with two convolutional layers followed by two bidirectional LSTM layers and a linear output layer.

#### Fluorescence

The fluorescence task is a protein engineering task which evaluates how fine-trained local genotypic changes can be captured to predict phenotypic expression, as measured by native fluorescence. The regression task is to predict the log-intensity of a mutant GFP sequence. Since this task is more sensitive to local than global information, we apply a mean-pool along the sequence dimension of the encoder output, *z*.

The data is from a Deep Mutational Scan (DMS) experiment from Sarkisyan et al. (2016), which measures fluorescence from derivative genotypes of the green fluorescent protein avGFP. Data splits are curated in Rao et al. (2019). Training and validation data are in a Hamming distance 3 neighborhood from the original protein, while the test data exhibits larger distribution shift and is from the Hamming distance 4-15 neighborhood.

For evaluation using a neural network head, `tape-proteins` uses the same MLP architecture as described in the remote homology task. The original benchmarks in Rao et al. (2019) compute an trainable attention-weighted mean prior to classification.

#### Stability

Stability is a protein engineering task which measures the most extreme concentration for which a protein can maintain its structure. This is a regression task to predict a stability score of proteins generated by *de novo* design. Since this task is also sensitive to fine-grained local effects, we use the mean along the encoder output *z* as a pooled embedding.

The data is from Rocklin et al. (2017), which measures the stability of proteins generated by parallel DNA synthesis, consisting of sequences from four protein topologies: *ααα*, *βαββ*, *αββα*, *ββαββ*. The stability score is the difference between the measured EC_50_ of the actual protein and its predicted EC_50_ in its unfolded state. Here, EC_50_ is the protease concentration at which 50% of cells pass the characterization threshold; note that it is measured on a log10 scale. Data splits are curated in Rao et al. (2019), such that the test set consists of seventeen 1-Hamming distance neighbourhoods from the training and validation datasets. A visualization of this test split is shown in Figure 4.

**Figure 4.**
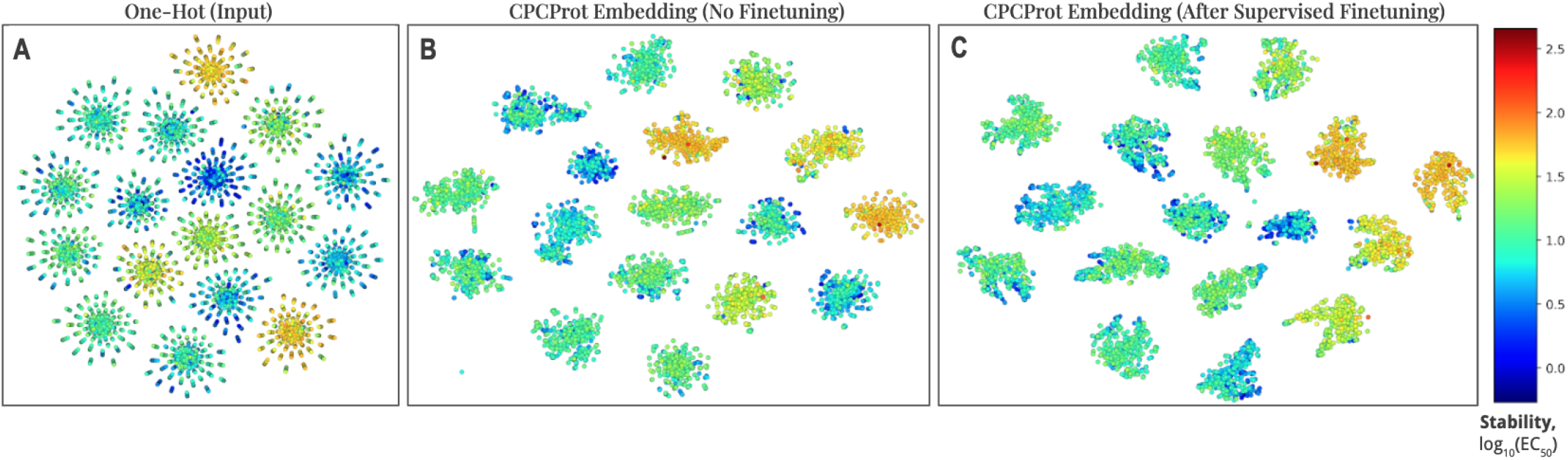
t-SNE visualization of proteins which are all 1-Hamming distance away from one of seventeen candidate proteins. Colors denote stability measurement on a log_10_ scale. The data corresponds to test set curated in the TAPE benchmarks (Rao et al., 2019) for the stability dataset from Rocklin et al. (2017).

For evaluation using a neural network head, as with remote homology and fluorescence, we use the provided MLP architecture, while the original benchmarks compute an trainable attention-weighted mean prior to classification.

## 5. Results

### 5.1. Downstream Tasks Evaluation

We evaluate the quality of the learned embeddings using the downstream tasks described in Section 4. As motivated in Section 4.2, we use different heads for downstream evaluation. In Table 1, we report results using neural network finetuning heads, and in Table 2–4, downstream results using simple linear and/or kNN models are reported.

**Table 1.** Embedding performance by downstream task using the default neural network finetuning head, compared against Tasks Assessing Protein Embeddings (TAPE) benchmarks (Rao et al., 2019). See Appendix C for dataset sizes by task.

|  | # of<br>Embedding<br>Parameters | Remote<br>Homology |  |  | Secondary<br>Structure |  |  | Stability | Fluorescence |
| --- | --- | --- | --- | --- | --- | --- | --- | --- | --- |
|  |  | Fold | Superfamily | Family | CB513 | CASP12 | TS115 |  |  |
| BERT | 92M | 0.21 | 0.34 | 0.88 | 0.73 | 0.71 | 0.77 | **0.73** | **0.68** |
| ResNet | 48M | 0.17 | 0.31 | 0.77 | **0.75** | **0.72** | **0.78** | **0.73** | 0.21 |
| LSTM | 44M | **0.26** | **0.43** | **0.92** | **0.75** | 0.70 | **0.78** | 0.69 | 0.67 |
| Bepler et al. | 19M | 0.17 | 0.20 | 0.79 | 0.73 | 0.70 | 0.76 | 0.64 | 0.33 |
| Unirep | 18M | 0.23 | 0.38 | 0.87 | 0.73 | **0.72** | 0.77 | **0.73** | 0.67 |
| One Hot | 0 | 0.09 | 0.08 | 0.39 | 0.69 | 0.68 | 0.72 | 0.19 | 0.14 |
| CPCProt | 1.7M | 0.12 | 0.12 | 0.48 | 0.69 | 0.70 | 0.73 | 0.65 | **0.68** |
| CPCProt<sub>GRU_large</sub> | 8.4M | 0.13 | 0.14 | 0.52 | 0.70 | 0.70 | 0.73 | 0.65 | **0.68** |
| CPCProt<sub>LSTM</sub> | 71M | 0.11 | 0.11 | 0.47 | 0.68 | 0.66 | 0.70 | 0.68 | **0.68** |

**Table 2.**
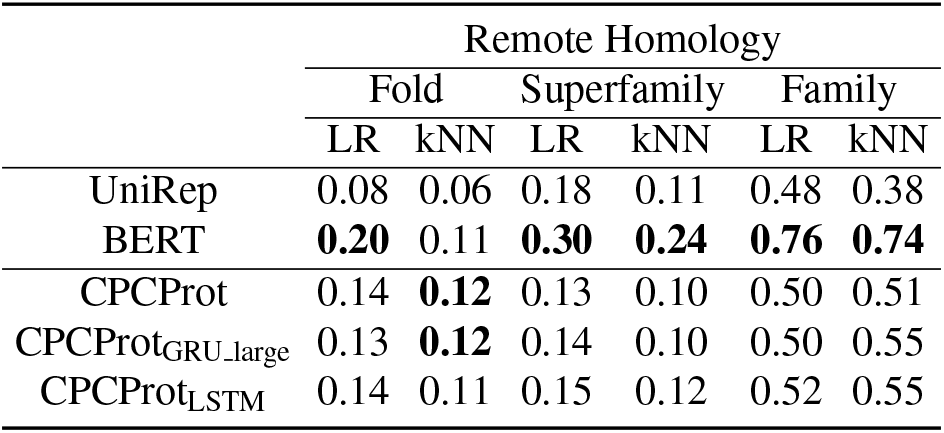
Downstream evaluation using logistic regression and kNN k-nearest-neighbours models (Top-1 accuracy).

|  | Remote Homology |  |  |  |  |  |
| --- | --- | --- | --- | --- | --- | --- |
|  | Fold |  | Superfamily |  | Family |  |
|  | LR | kNN | LR | kNN | LR | kNN |
| UniRep | 0.08 | 0.06 | 0.18 | 0.11 | 0.48 | 0.38 |
| BERT | **0.20** | 0.11 | **0.30** | **0.24** | **0.76** | **0.74** |
| CPCProt | 0.14 | **0.12** | 0.13 | 0.10 | 0.50 | 0.51 |
| CPCProt<sub>GRU_large</sub> | 0.13 | **0.12** | 0.14 | 0.10 | 0.50 | 0.55 |
| CPCProt<sub>LSTM</sub> | 0.14 | 0.11 | 0.15 | 0.12 | 0.52 | 0.55 |

We also evaluate two variant CPCProt models, selected during hyperparameter tuning by metrics which do not depend on downstream tasks, to avoid benchmark overfitting, and to examine the effect of increasing the number of parameters on downstream performance, as motivated in Section 4.1. The CPCProt_LSTM_ variant achieves the highest performance on the contrastive task, while the CPCProt_GRU_large_ variant achieves the highest performance on the 1NN Pfam family prediction task. Results on these tasks are reported in Appendix A. For all figures, however, we focus on our smallest 1.7M parameter CPCProt model.

#### CPCProt Performs Comparably with Baselines Using Fewer Parameters

CPCProt achieves comparable results as baselines on most tasks; however, we use only 2% of the number of embedding parameters of the largest bench-marked model, BERT (Rao et al., 2019) (Tables 1–5; Figure 1).

For the fluorescence task, CPCProt achieves higher *ρ* and lower MSE than other models for both a neural network fine-tuning head and most linear regression and kNN evaluations (Tables 1, 5; Appendix B). For the secondary structure and stability tasks, CPCProt achieves comparable performance with other neural network models, with a fraction of the number of parameters (Tables 1, 3, 4).

**Table 3.** Downstream evaluation using logistic regression models. Top-3 (Q3) accuracy is reported.

|  | Secondary Structure |  |  |
| --- | --- | --- | --- |
|  | CB513 | CASP12 | TS115 |
|  | LR | LR | LR |
| UniRep | 0.66 | 0.80 | 0.70 |
| BERT | **0.72** | **0.82** | **0.77** |
| CPCProt | 0.61 | 0.80 | 0.68 |
| CPCProt<sub>GRU_large</sub> | 0.62 | 0.80 | 0.69 |
| CPCProt<sub>LSTM</sub> | 0.62 | 0.80 | 0.69 |

**Table 4.** Downstream evaluation using linear regression and kNN models for the protein engineering task of stability (MSE and Spearman’s *ρ*).

|  | Stability |  |  |  |
| --- | --- | --- | --- | --- |
|  | LR |  | kNN |  |
| | MSE | $\rho$ | MSE | $\rho$ |
| UniRep | **0.21** | **0.62** | 0.24 | **0.57** |
| BERT | 0.36 | 0.39 | 0.23 | 0.49 |
| CPCProt | 0.34 | 0.55 | **0.18** | 0.51 |
| CPCProt<sub>GRU_large</sub> | 0.31 | **0.62** | **0.18** | 0.52 |
| CPCProt<sub>LSTM</sub> | 0.22 | **0.62** | 0.19 | 0.54 |

**Table 5.** Downstream evaluation using linear regression and kNN models for the protein engineering task of fluorescence (MSE and Spearman’s *ρ*).

|  | Fluorescence |  |  |  |
| --- | --- | --- | --- | --- |
|  | LR |  | kNN |  |
| | MSE | $\rho$ | MSE | $\rho$ |
| UniRep | 1.32 | 0.55 | **1.66** | 0.37 |
| BERT | 1.15 | 0.52 | 1.75 | 0.46 |
| CPCProt | 1.13 | 0.54 | 1.82 | 0.49 |
| CPCProt<sub>GRU_large</sub> | **0.81** | 0.63 | 1.84 | 0.50 |
| CPCProt<sub>LSTM</sub> | 0.85 | **0.67** | 1.80 | **0.51** |

For remote homology, when using a neural network for downstream evaluation, the difference in Top-1 accuracy between CPCProt and other neural network models is more drastic (Table 1); however, when using logistic regression and kNN for downstream classification, this gap in accuracy decreases (Table 2). As noted in Section 4.3, this is a highly class-imbalanced and multi-class task, and sometimes a one-shot problem for some classes in the Fold test set. This may contribute to the failure of all embeddings to generalize to the Fold test set even in the presence of labels, as well as the variability in changes to model performance when switching from a neural network to a simpler classifier (Tables 1, 2). As expected, as holdout test set distributions shift more from the training data, performance deteriorates for all models (Tables 1, 2).

#### Downstream Assessment is Inconsistent Using Different Models

For the purpose of using downstream tasks primarily as a means to evaluate the quality of embeddings, changing the downstream model used sometime result in preferring different models (Tables 1, 2, 3, 4, 5), as does using a different performance metric (i.e. MSE versus Spearman’s *ρ* for regression tasks) (Tables 4, 5, Appendix B). For example, though CPCProt appear to generalize poorly to the Fold test set for the remote homology task relative to baselines using a neural network classifier head (Table 1), it in fact outperforms both BERT and UniRep when using a kNN classifier (Table 2). For fluorescence and stability, which must capture fine-grained local effects, using a simple linear regression or kNN model seem to better differentiate performances of UniRep, BERT, and CPCProt variants (Tables 4, 5).

For stability and fluorescence, performance improves when using a finetuned neural network head for most embeddings, potentially reflecting the non-linear interactions needed to map local sequence effects to protein engineering measurements (Tables 1, 4, 5). In contrast, for secondary structure, finetuning with a higher capacity MLP actually *decreases* Q3 accuracy. For example, for the CASP12 test set, using a linear model instead of a MLP increases Q3 accuracy by 0.07 to 0.12 for BERT, UniRep, and CPCProt variants (Tables 1, 3). Note that the CASP12 test set consists only of 12 sequences, and the secondary structure training dataset is also the smallest out of the tasks compared (Appendix C).

### 5.2. CPCProt Qualitatively Captures Protein Stability Information without Supervised Training

As a motivating example to further “attribute responsibility” of downstream performance measurements to the pretrained model and finetuning procedures, we leverage the ability for t-SNE to preserve local neighbour information in high dimensions, and choose to examine the test set curated by Rao et al. (2019) of the stability data from Rocklin et al. (2017). The data was heldout to be from seventeen 1-Hamming distance neighbourhoods away from seventeen candidate proteins. Seventeen local structures cleanly emerge in the t-SNE embeddings from the raw one-hot encoded sequences alone (Figure 4). However, *within* each cluster of sequences derived from the same candidate protein, sequences close in stability are not assigned local neighbour relationships. After embedding using CPCProt, proteins close in stability measurements are also also close in t-SNE embedding space *within* each cluster, despite having never seen labels corresponding to stability during training. After updating embeddings via supervised end-to-end finetuning with stability labels using a MLP head, some new local structures emerge within clusters, though the ability for similarly stable proteins to cluster near each other do not improve drastically. This qualitatively suggests that the metrics presented in Table 1 can be reasonably attributed to the ability for CPCProt to capture implicit information about protein stability, despite not having seen any labels from stability experiments during training.

### 5.3. More Contrastive Learning Negative Samples Does Not Improve Performance

Theoretically, the InfoNCE estimator requires a large number of negative samples to reduce variance (Poole et al., 2019). Further, as formalized in Wang & Isola (2020), in the limit of infinite negative samples, the InfoMax objective has the desirable property of directly optimizing for the properties of alignment (similar samples are mapped to similar locations in the latent space) and uniformity (features preserve maximal information).

The standard practice, therefore, is that increasing the number of samples help performance, even at great computational costs. This has indeed yielded good results empirically (Hjelm et al., 2018; Tian et al., 2019; He et al., 2019; Bachman et al., 2019; Wu et al., 2018; Chen et al., 2020), However, in Saunshi et al. (2019), it is theoretically and empirically shown that larger number of negative samples may decrease downstream performance under certain conditions. As previous literature points out (Tschannen et al., 2019; Wang & Isola, 2020), this empirical and theoretical disjoint in how the number of negative samples affect performance render the success of using contrastive losses for representation learning more mysterious.

In Figure 5, we compare performance on the 1NN Pfam family prediction task (Section 4.1) for models pretrained using different numbers of negative samples. Our results corroborate the ideas in Saunshi et al. (2019) that down-stream performance does *not* empirically improve with the number of negative samples.

**Figure 5.**
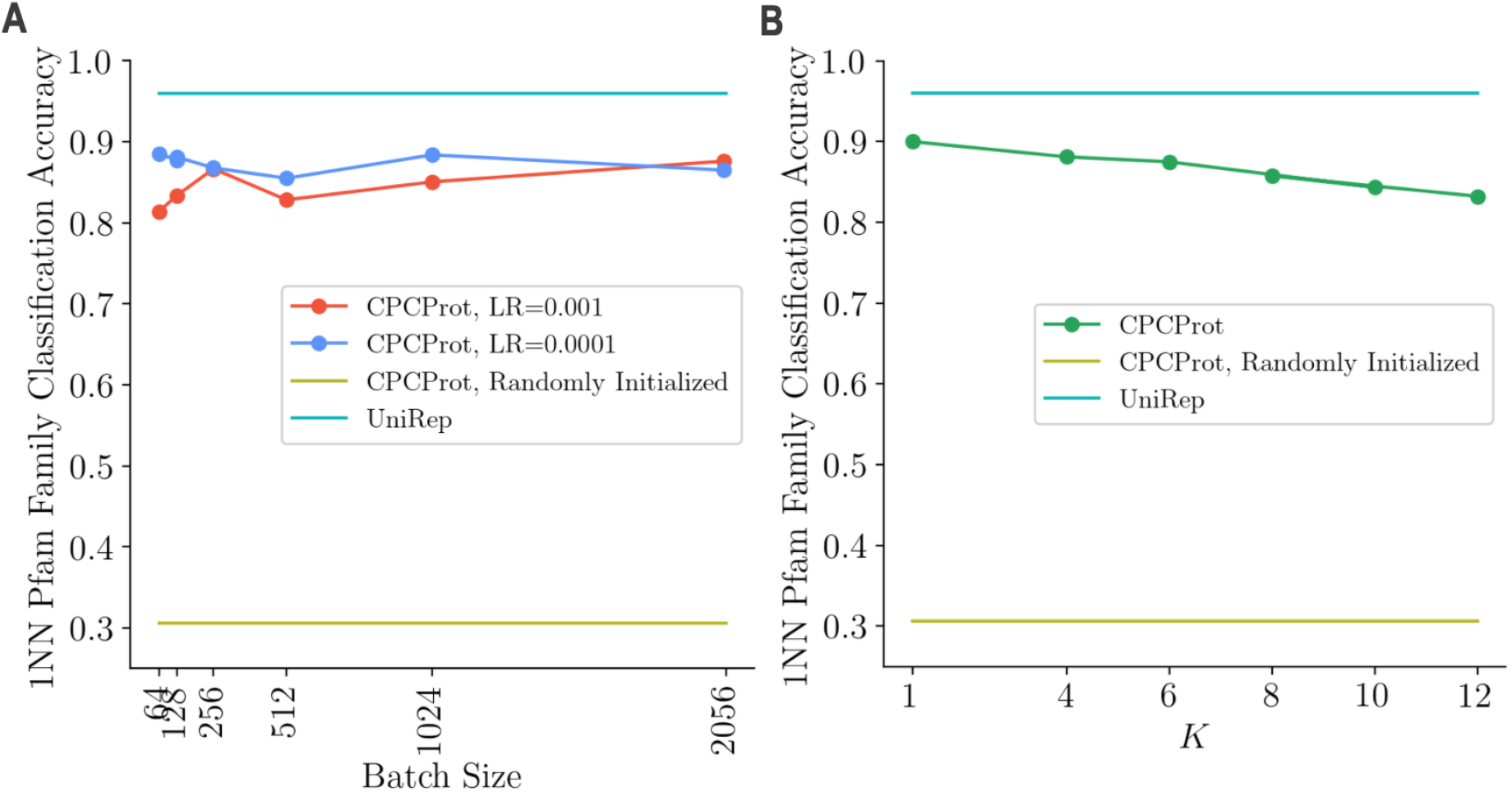
Downstream performance on a simple 1NN Pfam family classification task, using embedding models pretrained with different hyperparameters. For all plotted models, the default hyperparameters are LR=0.0001, batch size=1024, and K=4, using the same architecture as the 1.7M parameter CPCProt model. For reference, the performance of a randomly initialized CPCProt model and the 18M parameter UniRep model is also plotted. **(A)** Examination of the effect of batch size. Since we draw *N* − 1 negative samples in-batch, batch size is used as a proxy for the number of negative samples. **(B)** Examining the choice of *K*, that is, the number of “future” encoded amino acid patches which the model must learn to distinguish, using the context at that position.

We also further explore the effect of choosing *K*. A larger K necessitates that the model must learn information about amino acid patches further away given the context at position *t*. We train six different models with different settings of *K*, keeping all other hyperparameters consistent. On the 1NN Pfam family prediction task, asking the model to learn information about further away patches (i.e. larger *K*) decrease downstream performance. This may be in part due to the fact that a larger *K* increases the minimum sequence length needed, and results in more sequences discarded. For example, at *K* = 12, only Pfam sequences larger than 154 amino acids would be seen during training, which impacts performance when classifying shorter Pfam sequences.

### 5.4. Importance of Using a Per-*k* Critic

One major difference between CPC and other InfoMax methods is that the information maximization signal is formulated as an ordered autoregressive task (Hjelm et al., 2018), and uses a different critic for each position of *k*. In Figure 6, the accuracy on the contrastive task at each position of *t* + *k* for each *k* ∈ {1, 2, 3, 4} (averaged across all *t*) is visualized.

**Figure 6.**
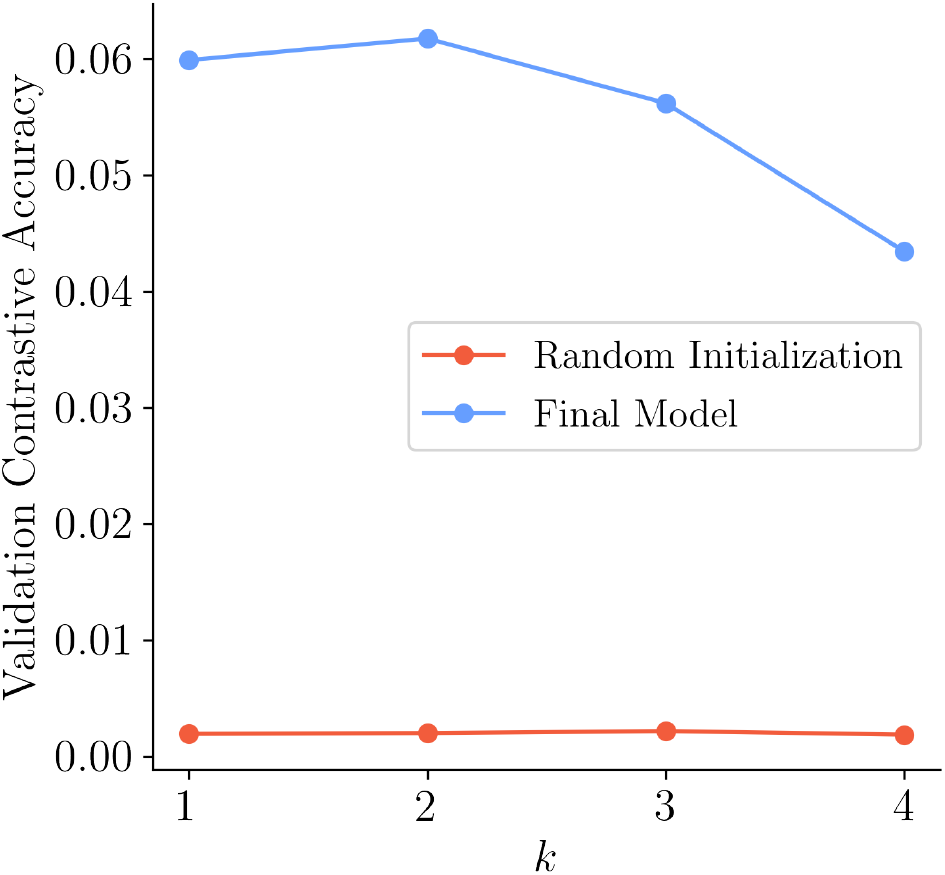
Comparison of accuracy on the contrastive task on the validation set, for a randomly initialized and the final 1.7M parameter CPCProt model, reported for each position of *t* + *k* (averaged over all *t* ∈ {*t*_*min*_, *t*_*min*_ + 1,…, *L*_*z*_ − *K*}). A random model achieves the expected accuracy of 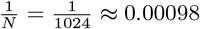, illustrating the difficulty of the pretraining task.

As expected, it becomes more difficult distinguish latent embeddings further away from the position from which the context is derived. The ability for the final model to solve this contrastive task is compared with a random model. This also illustrates the non-triviality of the contrastive task, which the model has learned to improve on after pretraining has converged. A validation batch size of 1024 is used, and the random model achieves roughly the expected accuracy of 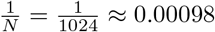.

## 6. Discussion

### Relationship Between Pretrained Model Size and Down-stream Performance

Table 1 and Figure 1 show that there is no clear connection between increasing the number of parameters (in the pretrained model only) and down-stream performance, contrary to the philosophy behind 567 million parameter NLP-inspired models for protein representations (Elnaggar et al., 2020). This is true even for variants of CPCProt, which were trained using the same self-supervised objective.

The finding that CPCProt achieves comparable results with less parameters may be a reflection of the overparameterization of existing protein embedding models in the field, or of a unique benefit conferred by the contrastive training. In any case, these results show that simply porting large models from NLP to proteins is not an efficient use of computational resources, and we encourage the community to further explore this relationship. As compared to currently-available protein embedding models, we note the suitability of CPCProt for downstream use-cases where model size is a key concern.

### Difficulties in Quantitatively Assessing Protein Embeddings

Though we explore using the pretraining objective (i.e. the contrastive accuracy) and a simple downstream task which assesses the capturing of global information (i.e. the 1NN Pfam family prediction task) to select hyperparameters for pretraining, we find that these performance metrics do not correlate with downstream performance for all tasks. This illustrates a difficulty of selecting pretraining hyperparameters for embeddings.

Furthermore, as explored in Section 5.1, even with the availability of downstream task benchmarks and data, it is difficult to quantitatively assess embedding performance, as results differ when using different models and performance metrics (i.e. MSE vs *ρ*). In some cases, where downstream accuracy is not the goal, it may be better use an embedding which outperforms other models when assessed by a linear downstream model and has presumably has captured more information, despite having a lower performance overall when compared to another model that provides better initialization for an neural network finetuning head.

Moreover, it is difficult to attribute quantitative performance on downstream tasks to the information captured by the embedding, or to the supervised finetuning procedures. We attempt to analyze this qualitatively in Figure 4. This is made more difficult by the inconsistency in whether if encoder weights should be frozen during training. While some works in contrastive learning freeze the encoder during fine-tuning (Wang & Isola, 2020), which makes sense as a means to directly evaluate the embeddings, large NLP embedding models such as BERT typically update parameters end-to-end (Devlin et al., 2018), as do protein models inspired by these NLP models (Rao et al., 2019), which makes sense as a means to evaluate the ability for these embeddings to achieve optimal performance for a specific task of interest.

Thus, we hope to highlight that downstream tasks should not be a definitive method to evaluate the goodness of an embedding. However, they may be good proxies to examine specific desiderata regarding global and local information or out-of-distribution generalization, when evaluated using a consistent protocol. Given the diversity of biological use cases, focusing on capturing crucial global information for one use case (e.g. long-range contacts) could be counter to another use case focused on highly localized effects (e.g. variant effect prediction). Quantitative metrics of down-stream performance should guide the choice of pretrained embeddings in a case-by-case manner.

### Limitations of CPC and In-Batch Negative Sampling

Though we apply the CPC method to proteins, there are other representation learning methods which fall under the InfoMax training scheme, which may also be applied to proteins. Further, the choice of negative samples to approximate the marginal distribution *p*(*z*′) affects the tightness of the bound (Tschannen et al., 2019; Poole et al., 2019; Saunshi et al., 2019). We choose negative samples in-batch for computational efficiency, but a more rigorous examination of the most apt negative sampling strategy for the protein modality is needed. As noted in Tschannen et al. (2019), maximizing MI does not always result in higher performance, and the empirical performance of recent InfoMax works may be due more to tuning the contrastive task and architecture. In fact, other works empirically demonstrate a U-shaped relationship between 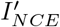 and downstream linear evaluation performance (Tian et al., 2020). The relationship between information maximization and good empirical results remains to be investigated, and we hope that our empirical examination of selecting the number of negative samples and the size of the local embedding window can extend beyond the protein modality.

## 7. Conclusion

In this work, we introduce CPCProt, which achieves comparable downstream performance as existing protein embedding models, at a fraction of the number of parameters. We further compare the effects of using different pretraining evaluation metrics and downstream models for evaluating embeddings on protein-related tasks, and find that there is poor consistency in how models compare against one another, illustrating the difficulty in defining the “goodness” of an embedding for biological use cases. In addition, we empirically evaluate hyperparameters affecting contrastive training for representation learning, and empirically corroborate the theoretical result that additional negative samples do not always improve performance (Saunshi et al., 2019). In academic and industry settings where model size is a key concern, CPCProt may be a desirable embedding method as compared to other currently available methods, as it fits easily on a single GPU, and achieves results in the same neighborhood of performance as larger pretrained models. We hope that this work can inform the development of other embedding models for biological sequences, as well as contrastive representation learning methods beyond the protein sequence modality.

## Acknowledgements

Amy X. Lu was funded by the NSERC Canada Graduate Scholarships-Master’s award. Alan M. Moses holds a Tier II Canada Research Chair. Maryzeh Ghassemi is funded in part by Microsoft Research, a CIFAR AI Chair at the Vector Institute, a Canada Research Council Chair, and an NSERC Discovery Grant. Computing resources for this work was provided by the Vector Institute. The authors thank Nick Bhattacharya, Alex X. Lu, Adam Riesselman, Denny Wu, and Kevin Yang for insightful input.

## A. Evaluation Results for Pretraining Hyperparameter Selection

**Table 6.** Results on evaluation metrics for selecting hyperparameters and architectures. The contrastive accuracy is the NCE task, using a batch size of 1024, such that a random model achieves an expected accuracy of 0.00098. Our CPCProt_LSTM_ variant achieves the highest contrastive accuracy of all evaluated models, while the CPCProt_GRU_large_ variant achieves the highest accuracy on the 1NN Pfam family prediction task. We report models selected by metrics which do not depend on downstream tasks to avoid benchmark overfitting, and to examine the effect of increasing the number of parameters on desired properties of embeddings. For reference, UniRep achieves 95% accuracy on our 1NN Pfam family prediction task and dataset.

|  | Pretraining Validation Dataset |  | Downstream Tasks Validation Dataset |  |  |  |
| --- | --- | --- | --- | --- | --- | --- |
|  | Contrastive Accuracy | 1NN | Fluorescence | Stability | Remote Homology | Secondary Structure |
| CPCProt | 0.04 | 0.89 | **0.80** | **0.68** | 0.11 | **0.71** |
| CPCProt<sub>LSTM</sub> | **0.09** | 0.91 | 0.79 | 0.68 | **0.12** | 0.66 |
| CPCProt<sub>GRU_large</sub> | 0.05 | **0.93** | 0.72 | 0.67 | 0.10 | 0.69 |

## B. Mean-Squared Error on Fluorescence Task, MLP Evaluation Protocol

**Table 7.** Mean-squared error of various embedding methods on the fluorescence task.

|  | MSE |
| --- | --- |
| BERT | 0.22 |
| Unirep | 0.20 |
| ResNet | 3.04 |
| LSTM | 0.19 |
| One Hot | 2.69 |
| CPCProt | 0.14 |
| CPCProt<sub>LSTM</sub> | **0.10** |
| CPCProt<sub>GRU_large</sub> | **0.10** |

## C. Downstream Tasks Dataset Sizes

|  |  | **Test Dataset Size** | **Training Dataset Size** |
| --- | --- | --- | --- |
| **Remote Homology** | Fold | 718 | 12312 |
|  | Superfamily | 1254 |  |
|  | Family | 1272 |  |
| **Secondary Structure** | CB513 | 513 | 8678 |
|  | CASP12 | 21 |  |
|  | TS115 | 115 |  |
| **Stability** |  | 12851 | 53614 |
| **Fluorescence** |  | 27217 | 21446 |

## Footnotes

1 In this work, we use the more general term “neural network finetuning” as opposed to “MLP finetuning” in the contrastive representation learning literature, as one of the protein-related downstream tasks is a sequence-to-sequence task which uses a CNN head.

## References

Abriata, L. A., Tamò, G. E., Monastyrskyy, B., Kryshtafovych, A., and Dal Peraro, M. Assessment of hard target modeling in casp12 reveals an emerging role of alignment-based contact prediction methods. Proteins: Structure, Function, and Bioinformatics, 86:97–112, 2018.

Adami, C. Information theory in molecular biology. Physics of Life Reviews, 1(1):3–22, 2004.

Alemi, A. A., Fischer, I., Dillon, J. V., and Murphy, K. Deep variational information bottleneck. arXiv preprint arXiv:1612.00410, 2016.

Alley, E. C., Khimulya, G., Biswas, S., AlQuraishi, M., and Church, G. M. Unified rational protein engineering with sequence-only deep representation learning. bioRxiv, pp. 589333, 2019.

Armenteros, J. J. A., Johansen, A. R., Winther, O., and Nielsen, H. Language modelling for biological sequences–curated datasets and baselines. 2019.

Bachman, P., Hjelm, R. D., and Buchwalter, W. Learning representations by maximizing mutual information across views. In Advances in Neural Information Processing Systems, pp. 15509–15519, 2019.

Belghazi, M. I., Baratin, A., Rajeswar, S., Ozair, S., Bengio, Y., Courville, A., and Hjelm, R. D. Mine: mutual information neural estimation. arXiv preprint arXiv:1801.04062, 2018.

Bepler, T. and Berger, B. Learning protein sequence embeddings using information from structure. arXiv preprint arXiv:1902.08661, 2019.

Chen, T., Kornblith, S., Norouzi, M., and Hinton, G. A simple framework for contrastive learning of visual representations. arXiv preprint arXiv:2002.05709, 2020.

Consortium, U. Uniprot: a worldwide hub of protein knowledge. Nucleic acids research, 47(D1):D506–D515, 2019.

Cuff, J. A. and Barton, G. J. Evaluation and improvement of multiple sequence methods for protein secondary structure prediction. Proteins: Structure, Function, and Bioinformatics, 34(4):508–519, 1999.

Devlin, J., Chang, M.-W., Lee, K., and Toutanova, K. Bert: Pre-training of deep bidirectional transformers for language understanding. arXiv preprint arXiv:1810.04805, 2018.

Donsker, M. D. and Varadhan, S. S. Asymptotic evaluation of certain markov process expectations for large time. iv. Communications on Pure and Applied Mathematics, 36 (2):183–212, 1983.

El-Gebali, S., Mistry, J., Bateman, A., Eddy, S. R., Luciani, A., Potter, S. C., Qureshi, M., Richardson, L. J., Salazar, G. A., Smart, A., Sonnhammer, E. L. L., Hirsh, L., Paladin, L., Piovesan, D., Tosatto, S. C. E., and Finn, R. D. The Pfam protein families database in 2019. Nucleic Acids Research, 47(D1): D427–D432, 2019. ISSN 0305-1048. doi: 10.1093/nar/gky995. URL https://academic.oup.com/nar/article/47/D1/D427/5144153.

Elnaggar, A., Heinzinger, M., Dallago, C., and Rost, B. End-to-end multitask learning, from protein language to protein features without alignments. bioRxiv, pp. 864405, 2019.

Elnaggar, A., Heinzinger, M., Dallago, C., Rihawi, G., Wang, Y., Jones, L., Gibbs, T., Feher, T., Angerer, C., Bhowmik, D., et al. Prottrans: Towards cracking the language of life’s code through self-supervised deep learning and high performance computing. arXiv preprint arXiv:2007.06225, 2020.

Fox, N. K., Brenner, S. E., and Chandonia, J.-M. Scope: Structural classification of proteins—extended, integrating scop and astral data and classification of new structures. Nucleic acids research, 42(D1):D304–D309, 2013.

Gatlin, L. L. et al. Information theory and the living system. Columbia University Press, 1972.

Gligorijevic, V., Renfrew, P. D., Kosciolek, T., Leman, J. K., Cho, K., Vatanen, T., Berenberg, D., Taylor, B. C., Fisk, I. M., Xavier, R. J., et al. Structure-based function prediction using graph convolutional networks. bioRxiv, pp. 786236, 2019.

Gutmann, M. and Hyvärinen, A. Noise-contrastive estimation: A new estimation principle for unnormalized statistical models. In Proceedings of the Thirteenth International Conference on Artificial Intelligence and Statistics, pp. 297–304, 2010.

He, K., Fan, H., Wu, Y., Xie, S., and Girshick, R. Momentum contrast for unsupervised visual representation learning. arXiv preprint arXiv:1911.05722, 2019.

Heinzinger, M., Elnaggar, A., Wang, Y., Dallago, C., Nechaev, D., Matthes, F., and Rost, B. Modeling the language of life-deep learning protein sequences. bioRxiv, pp. 614313, 2019.

Hénaff, O. J., Razavi, A., Doersch, C., Eslami, S., and Oord, A. v. d. Data-efficient image recognition with contrastive predictive coding. arXiv preprint arXiv:1905.09272, 2019.

Hjelm, R. D., Fedorov, A., Lavoie-Marchildon, S., Grewal, K., Bachman, P., Trischler, A., and Bengio, Y. Learning deep representations by mutual information estimation and maximization. arXiv preprint arXiv:1808.06670, 2018.

Hou, J., Adhikari, B., and Cheng, J. Deepsf: deep convolutional neural network for mapping protein sequences to folds. Bioinformatics, 34(8):1295–1303, 2018.

Kabsch, W. and Sander, C. Dictionary of protein secondary structure: pattern recognition of hydrogen-bonded and geometrical features. Biopolymers: Original Research on Biomolecules, 22(12):2577–2637, 1983.

Kingma, D. P. and Ba, J. Adam: A method for stochastic optimization. arXiv preprint arXiv:1412.6980, 2014.

Kingma, D. P. and Welling, M. Auto-encoding variational bayes. arXiv preprint arXiv:1312.6114, 2013.

Klausen, M. S., Jespersen, M. C., Nielsen, H., Jensen, K. K., Jurtz, V. I., Soenderby, C. K., Sommer, M. O. A., Winther, O., Nielsen, M., Petersen, B., et al. Netsurfp-2.0: Improved prediction of protein structural features by integrated deep learning. Proteins: Structure, Function, and Bioinformatics, 2019.

Kong, L., d’Autume, C. d. M., Ling, W., Yu, L., Dai, Z., and Yogatama, D. A mutual information maximization perspective of language representation learning. arXiv preprint arXiv:1910.08350, 2019.

Korber, B. T., Farber, R. M., Wolpert, D. H., and Lapedes, A. S. Covariation of mutations in the v3 loop of human immunodeficiency virus type 1 envelope protein: an information theoretic analysis. Proceedings of the National Academy of Sciences, 90(15):7176–7180, 1993.

Linsker, R. Self-organization in a perceptual network. Computer, 21(3):105–117, 1988.

Löwe, S., O’Connor, P., and Veeling, B. Putting an end to end-to-end: Gradient-isolated learning of representations. In Advances in Neural Information Processing Systems, pp. 3033–3045, 2019.

Madani, A., McCann, B., Naik, N., Keskar, N. S., Anand, N., Eguchi, R. R., Huang, P.-S., and Socher, R. Progen: Language modeling for protein generation. arXiv preprint arXiv:2004.03497, 2020.

Moult, J., Fidelis, K., Kryshtafovych, A., Schwede, T., and Tramontano, A. Critical assessment of methods of protein structure prediction (CASP)-Round XII. Proteins: Structure, Function, and Bioinformatics, 86:7–15, 2018. ISSN 08873585. doi: 10.1002/prot.25415. URL http://doi.wiley.com/10.1002/prot.25415.

Nguyen, X., Wainwright, M. J., and Jordan, M. I. Estimating divergence functionals and the likelihood ratio by convex risk minimization. IEEE Transactions on Information Theory, 56(11):5847–5861, 2010.

Nourani, E., Asgari, E., McHardy, A. C., and Mofrad, M. R. Tripletprot: Deep representation learning of proteins based on siamese networks. bioRxiv, 2020.

Oord, A. v. d., Li, Y., and Vinyals, O. Representation learning with contrastive predictive coding. arXiv preprint arXiv:1807.03748, 2018.

Pathak, D., Krahenbuhl, P., Donahue, J., Darrell, T., and Efros, A. A. Context encoders: Feature learning by inpainting. In Proceedings of the IEEE conference on computer vision and pattern recognition, pp. 2536–2544, 2016.

Poole, B., Ozair, S., Oord, A. v. d., Alemi, A. A., and Tucker, G. On variational bounds of mutual information. arXiv preprint arXiv:1905.06922, 2019.

Pritišanac, I., Vernon, R. M., Moses, A. M., and For-man Kay, J. D. Entropy and information within intrinsically disordered protein regions. Entropy, 21(7):662, 2019.

Rao, G. S., Hamid, Z., and Rao, J. S. The information content of dna and evolution. Journal of theoretical biology, 81(4):803–807, 1979.

Rao, R., Bhattacharya, N., Thomas, N., Duan, Y., Chen, P., Canny, J., Abbeel, P., and Song, Y. Evaluating protein transfer learning with tape. In Advances in Neural Information Processing Systems, pp. 9686–9698, 2019.

Recht, B., Roelofs, R., Schmidt, L., and Shankar, V. Do imagenet classifiers generalize to imagenet? arXiv preprint arXiv:1902.10811, 2019.

Riesselman, A. J., Shin, J.-E., Kollasch, A. W., McMahon, C., Simon, E., Sander, C., Manglik, A., Kruse, A. C., and Marks, D. S. Accelerating protein design using autoregressive generative models. bioRxiv, pp. 757252, 2019.

Rives, A., Goyal, S., Meier, J., Guo, D., Ott, M., Zitnick, C. L., Ma, J., and Fergus, R. Biological structure and function emerge from scaling unsupervised learning to 250 million protein sequences. bioRxiv, pp. 622803, 2019.

Rivière, M., Joulin, A., Mazaré, P.-E., and Dupoux, E. Unsupervised pretraining transfers well across languages. arXiv preprint arXiv:2002.02848, 2020.

Rocklin, G. J., Chidyausiku, T. M., Goreshnik, I., Ford, A., Houliston, S., Lemak, A., Carter, L., Ravichandran, R., Mulligan, V. K., Chevalier, A., et al. Global analysis of protein folding using massively parallel design, synthesis, and testing. Science, 357(6347):168–175, 2017.

Roman-Roldan, R., Bernaola-Galvan, P., and Oliver, J. Application of information theory to dna sequence analysis: a review. Pattern recognition, 29(7):1187–1194, 1996.

Sarkisyan, K. S., Bolotin, D. A., Meer, M. V., Usmanova, D. R., Mishin, A. S., Sharonov, G. V., Ivankov, D. N., Bozhanova, N. G., Baranov, M. S., Soylemez, O., et al. Local fitness landscape of the green fluorescent protein. Nature, 533(7603):397, 2016.

Saunshi, N., Plevrakis, O., Arora, S., Khodak, M., and Khandeparkar, H. A theoretical analysis of contrastive unsupervised representation learning. In International Conference on Machine Learning, pp. 5628–5637, 2019.

Schneider, T. D. and Stephens, R. M. Sequence logos: a new way to display consensus sequences. Nucleic acids research, 18(20):6097–6100, 1990.

Stormo, G. D. Dna binding sites: representation and discovery. Bioinformatics, 16(1):16–23, 2000.

Tian, Y., Krishnan, D., and Isola, P. Contrastive multiview coding. arXiv preprint arXiv:1906.05849, 2019.

Tian, Y., Sun, C., Poole, B., Krishnan, D., Schmid, C., and Isola, P. What makes for good views for contrastive learning. arXiv preprint arXiv:2005.10243, 2020.

Tschannen, M., Djolonga, J., Rubenstein, P. K., Gelly, S., and Lucic, M. On mutual information maximization for representation learning. arXiv preprint arXiv:1907.13625, 2019.

Vinga, S. Information theory applications for biological sequence analysis. Briefings in bioinformatics, 15(3): 376–389, 2014.

Wang, T. and Isola, P. Understanding contrastive representation learning through alignment and uniformity on the hypersphere. arXiv preprint arXiv:2005.10242, 2020.

Wu, Z., Xiong, Y., Yu, S. X., and Lin, D. Unsupervised feature learning via non-parametric instance discrimination. In Proceedings of the IEEE Conference on Computer Vision and Pattern Recognition, pp. 3733–3742, 2018.

Yang, K. K., Wu, Z., Bedbrook, C. N., and Arnold, F. H. Learned protein embeddings for machine learning. Bioinformatics, 34(15):2642–2648, 2018a.

Yang, Y., Gao, J., Wang, J., Heffernan, R., Hanson, J., Paliwal, K., and Zhou, Y. Sixty-five years of the long march in protein secondary structure prediction: the final stretch? Briefings in bioinformatics, 19(3):482–494, 2018b.

